# Human brain cell-type-specific aging clocks based on single-nuclei transcriptomics

**DOI:** 10.1101/2025.02.28.640749

**Authors:** Chandramouli Muralidharan, Enikő Zakar-Polyák, Anita Adami, Anna A. Abbas, Yogita Sharma, Raquel Garza, Jenny G. Johansson, Diahann A. M. Atacho, Éva Renner, Miklós Palkovits, Csaba Kerepesi, Johan Jakobsson, Karolina Pircs

## Abstract

Aging is the primary risk factor for most neurodegenerative diseases, yet the cell-type-specific progression of brain aging remains poorly understood. Here, we developed human cell-type-specific transcriptomic aging clocks using high-quality single-nucleus RNA sequencing data from post mortem human prefrontal cortex tissue of 31 donors aged 18 – 94 years, encompassing 73,941 high-quality nuclei. We observed distinct transcriptomic changes across major cell types, including upregulation of inflammatory response genes in microglia from older samples. Aging clocks trained on each major cell type accurately predicted chronological age and remained robust in independent single-nucleus RNA-sequencing datasets, underscoring their broad applicability. These findings demonstrate the feasibility of cell-type-specific transcriptomic clocks to measure biological aging in the human brain and highlight potential mechanisms of selective vulnerability in neurodegenerative diseases. We anticipate these clocks will serve as a basis for further studies in other brain regions and more diverse populations, ultimately advancing our understanding of age-related neurodegenerative processes at the single-cell level.

## INTRODUCTION

Aging involves complex molecular changes that gradually lead to the functional decline of cells and organs, including the different cell types in the brain^1,2^. Aging is a major risk factor for most neurodegenerative diseases, including Alzheimer’s and Parkinson’s disease^3^. However, our understanding of how different cell types age in the brain, the molecular mechanisms underlying their aging processes and how these changes contribute to disease pathophysiology remains limited. To better understand these processes, it will be important to develop tools that accurately measure molecular age-related changes in the different cell types in the brain and how these processes are altered in disease.

In recent years, several molecular aging clocks have been developed that use different types of omics data in combination with machine learning algorithms to predict biological age^4–9^. These tools offer a quantitative measure of aging beyond chronological age, enabling the identification of individuals or tissues at risk for accelerated aging and providing insights into underlying molecular pathways. With few exceptions^4,10–13^, these aging clocks are derived from bulk tissue^6–8,14,15^ and therefore lack cell-type-specific resolution. With an increasing number of studies pointing to cell-type-specific mechanisms of aging in the human brain^16–20^, such bulk-trained clocks have a major limitation in accurately capturing age-related changes. In a recent study in the mouse brain, a cell-type-specific transcriptomic aging clock based on single-cell RNA sequencing was developed. It not only accurately predicted the chronological age for different cell types but also modelled the reversal of aging with exercise and rejuvenation^4^. This highlights the need to develop human brain cell-type-specific aging clocks that can predict the age of individual cells from different cell types. Such a tool would have the potential to identify the selective vulnerability of particular cell types in human age-related neurodegenerative disorders.

In this study, we performed single-nuclei RNA sequencing on 31 post mortem human prefrontal cortex tissues from young, middle-aged and old donors. Using this dataset, we found cell-type-specific age-related transcriptomic differences, allowing us to develop a human single-nuclei-based transcriptomic aging clock for different cell types in the brain, which could be validated using publicly available single nuclei RNA-sequencing datasets.

## RESULTS

### Single-nuclei RNA sequencing of a human aging cohort

To study cell-type-specific changes during human brain aging, we performed single-nuclei RNA sequencing (snRNA-seq) on post mortem human prefrontal cortex tissue derived from 31 donors aged between 18 and 94 years at death (Fig. 1a-d, Table 1 and Supplementary Data 1a). We sequenced fresh frozen punch biopsies originating from the ventrolateral prefrontal cortex or the middle frontal gyrus, two brain regions that are a part of the prefrontal cortex, which has been implicated in age-related neurodegenerative disorders such as Alzheimer’s disease^21–24^. Tissue samples were distributed across three age groups, including tissue coming from young (18 – 39 years), middle-aged (40 – 59 years) and old (60 + years) individuals (Fig. 1b). Tissue was collected with a short median post mortem interval (PMI) of 4.5 (2 – 12) hours (Fig. 1c) and was obtained from both sexes (Fig. 1d). After sequencing and quality control we obtained data from 73,941 high quality nuclei (Supplementary Fig. 1a-d and Supplementary Data 1b). We performed an unbiased clustering resulting in 19 distinct cell clusters (Fig. 1e). Using the expression of cell-type-specific marker genes we identified all major cell types present in the human prefrontal cortex, including oligodendrocytes, astrocytes, oligodendrocyte progenitor cells (OPCs), microglia, inhibitory neurons, and excitatory neurons (Fig. 1f-g, Supplementary Fig. 1e and Supplementary Data 2). All major cell types were represented across all samples (Fig. 1h, Supplementary Fig. 1f, and Supplementary Data 1c) and across the different age groups (Fig. 1j). In addition, these cell types were the most abundant in the dataset (Fig. 1i), making them suitable for comparison.

**Figure 1.**
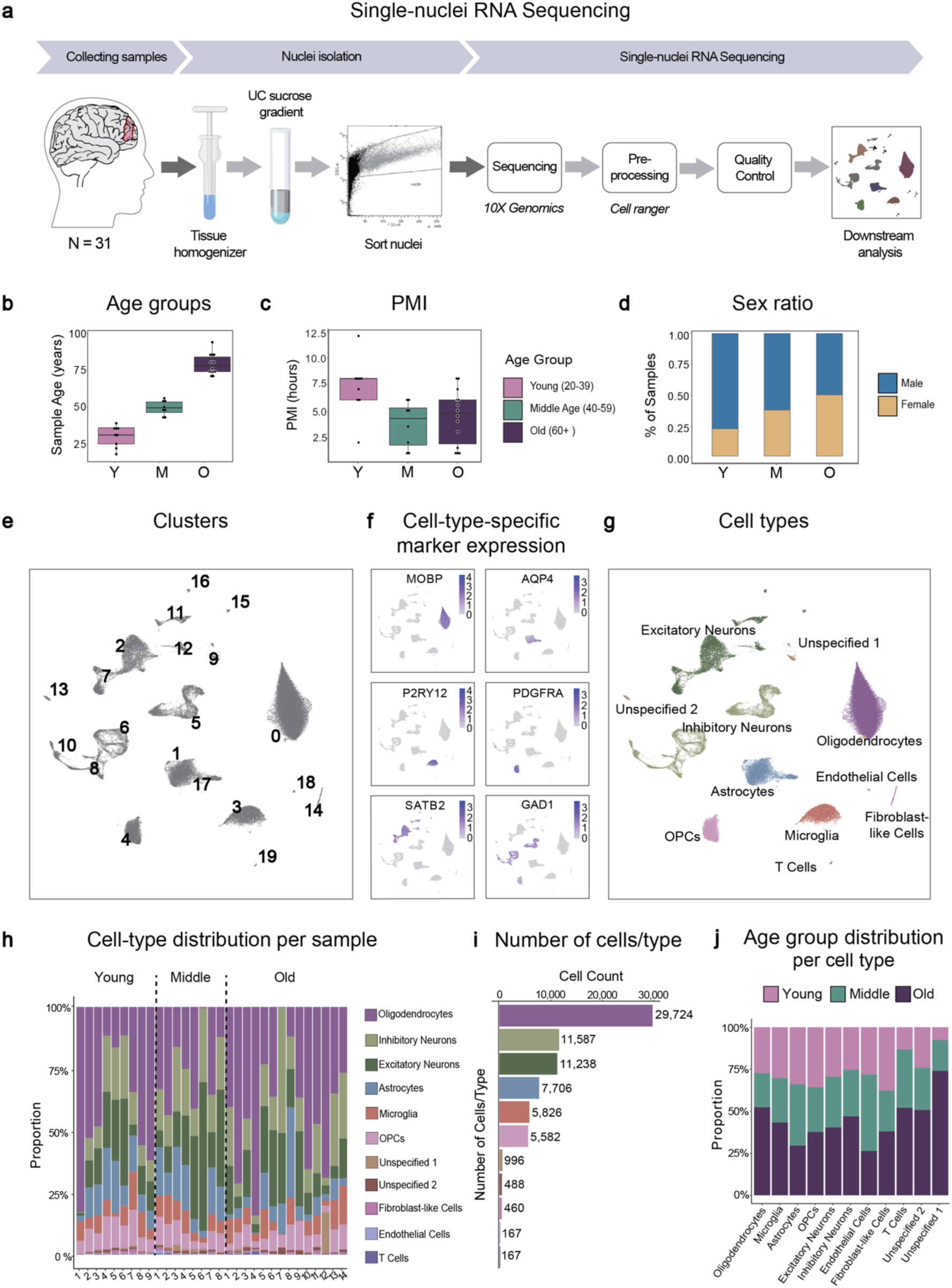
Single-nucleus RNA sequencing of human post mortem prefrontal cortex tissue from an aging cohort. (a) Schematic of the experimental workflow for snRNA-seq. Nuclei were isolated and sorted from frozen human post mortem prefrontal cortex tissue samples from 31 donors, followed by snRNA-seq and downstream analysis. (b) Box plot showing the age distribution of samples in the different age groups. Samples in the young age group (Y, n = 8), range 18 - 39 years, 40 - 59 years in the middle-aged group (M, n = 7) and 60 - 94 years in the old age group (O, n = 14). (c) Box plot showing the distribution of the post mortem interval (PMI) of samples in the different age groups. (d) Bar chart showing the proportion of the samples of each sex in the different age groups. (e) UMAP plot annotated with the 19 identified cell clusters. (f) Projection of gene expression of canonical markers of different cell types in the adult human prefrontal cortex. (g) UMAP plot annotated with the identified cell types. (h) Bar plot showing the proportion of cells of different cell types in all the samples. (i) Bar graph showing the number of cells in the dataset from each identified cell type. (j) Bar graph showing the proportion of cells of different age groups within each cell type. See also Supplementary Figure 1.

**Table 1:**
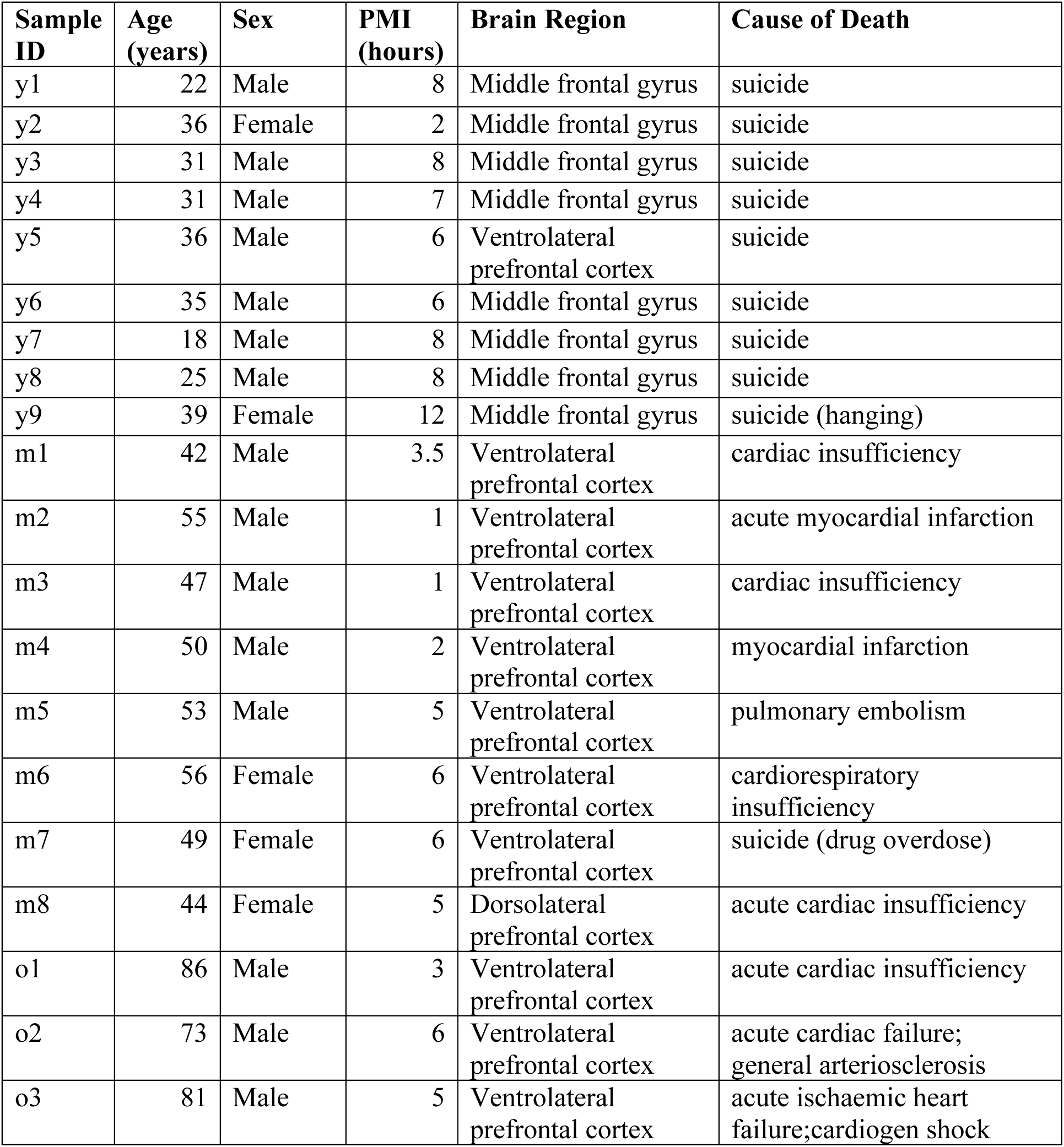
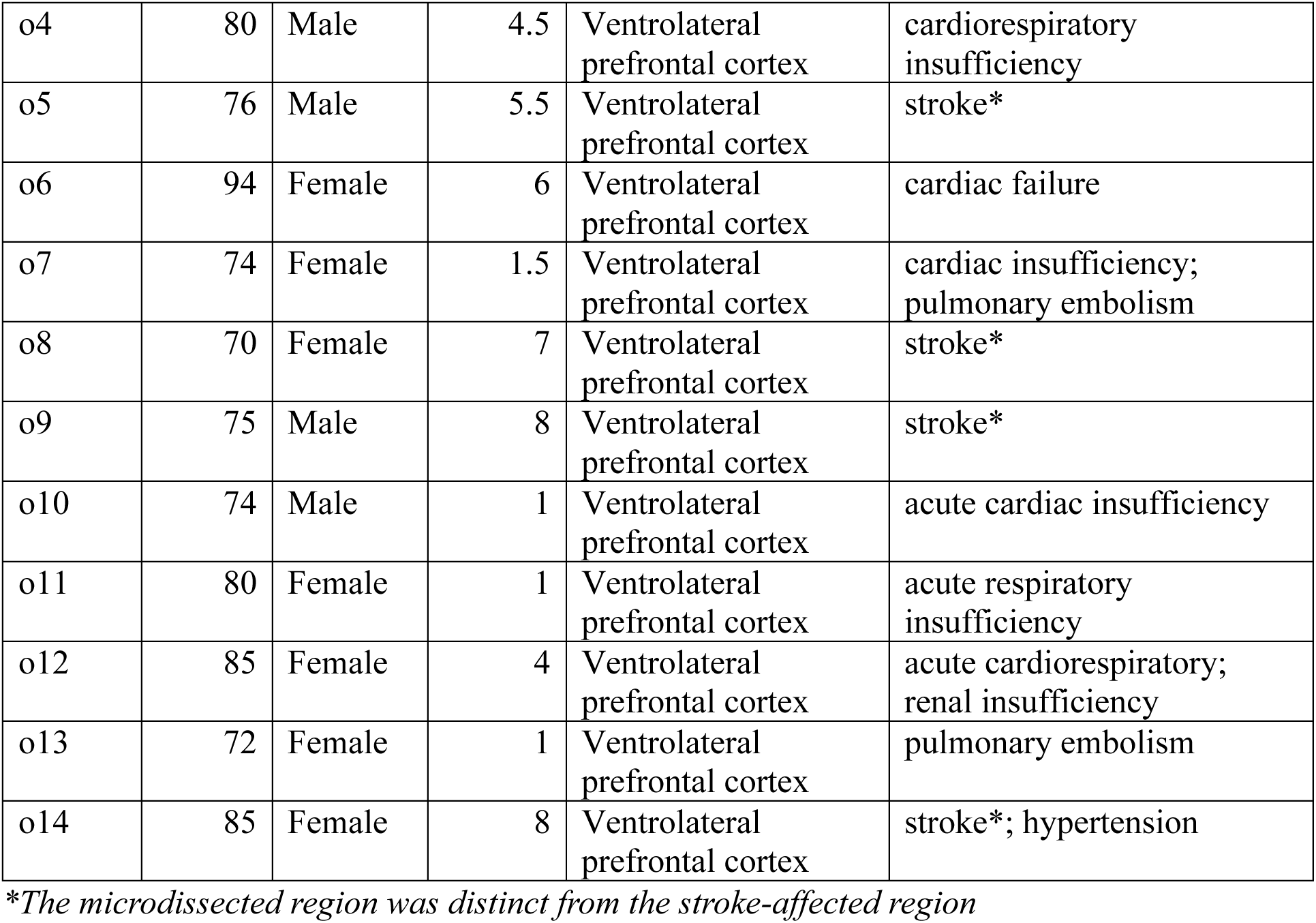
Demographic and clinical data of the human prefrontal cortical samples.

### Age-Related Transcriptomic Changes in Microglia and Astrocytes

First, we investigated cell-type-specific age-related changes in our dataset by performing differential gene expression analysis between different age groups. We found distinct age-related transcriptomic changes in each of the different cell types (Fig. 2a, d, Supplementary Fig. 2a, c, d, f and Supplementary Data 3). For example, in microglia, we observed inflammatory response genes significantly upregulated in the old age group when compared to the young or the middle age groups, such as *FOXP1*, *TLR2* and *CD163*, while homeostatic microglial markers, such as *CX3CR1*, *P2RY12* and *P2RY13*, were downregulated (Fig. 2b and Supplementary Data 3g-l). Gene over-representation test of the differentially expressed genes revealed an enrichment of terms related to inflammatory response (Fig. 2c Supplementary Figure 3a and Supplementary Data 4d-f). These observations are consistent with previous studies on microglial aging, which reported an increased inflammatory response during aging^25,26^. In astrocytes, we found an upregulation of genes associated with reactive astrogliosis, such as *TPST1*, *SAMD4A*, *CNN3*, *STAT3* and *SORBS1* (Fig. 2e and Supplementary Data 3m-r), in the old group when compared to young or middle-aged groups. This is in line with previous studies showing that reactive astrogliosis occurs during inflammation in the aging brain^27–29^. Gene ontology analysis revealed that terms related to protein misfolding, a hallmark of brain aging^1,2^ were also enriched in differentially expressed genes in astrocytes (Fig. 2f and Supplementary Data 4g-h). In oligodendrocytes, gene ontology analysis revealed chaperone-mediated protein-folding term to be enriched (Supplementary Fig. 2b and Supplementary Data 4a-c). Excitatory neurons revealed gene ontology terms related to protein translation to be enriched (Supplementary Fig. 2e and Supplementary Data 4j-l), while in inhibitory neurons and OPCs, no significant terms were enriched in old vs young age group comparisons.

**Figure 2.**
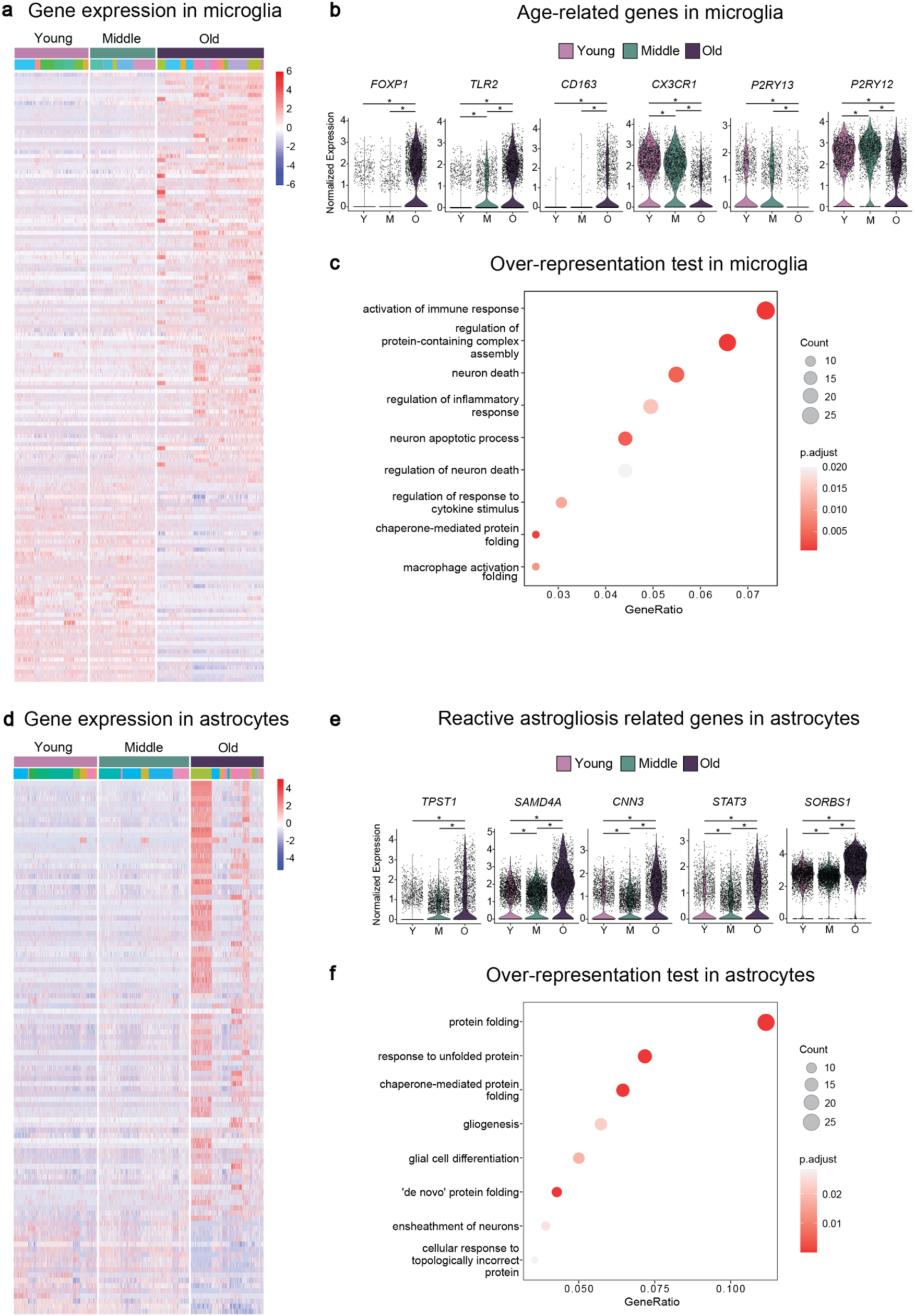
Age-related transcriptomic changes are present in microglia and astrocytes. (a) Heatmap showing the expression of the top differentially expressed genes (-0.75 > average log2 fold-change > 0.75, adj. p-value < 0.05) in microglia when comparing old vs young age groups. Different colours below the young, middle and old ages bar represents cells originating from different individuals. (b) Violin plots showing the expression of several previously identified microglial aging genes. (c) Plot showing selected gene ontology terms enriched from genes differentially expressed in old microglia. (d) Heatmap showing the expression of the top differentially expressed genes (-0.75 > average log2 fold-change > 0.75, adj. p-value < 0.05) in astrocytes when comparing old vs young age groups. Different colours below the young, middle and old ages bar represents cells originating from different individuals. (e) Violin plots showing the expression of selected reactive astrogliosis genes. (f) Plots showing selected gene ontology terms enriched from genes differentially expressed in old astrocytes. \**P* < 0.05; Wilcoxon rank sum test was used for each comparison in b and e, while ClusterProfiler’s gene-overrepresentation analysis along with Bonferroni-Hochberg correction for multiplicity was used in e and f. See also Supplementary Figure 2.

In summary, these results demonstrate that age-related transcriptomic changes are present and can be detected in our snRNA-seq dataset. Importantly, this transcriptional response to aging is different in each cell type, highlighting the relevance of the development of cell-type-specific transcriptional clock algorithms using this dataset.

### Development and evaluation of cell-type-specific transcriptomic aging clocks for the human brain

To measure the extent of aging in each cell type based on transcriptomic information, we trained ElasticNet regression models^30,31^ on our dataset via fivefold cross-validation using genes as features and age as the target. We focused on the most abundant cell types in the dataset, including oligodendrocytes, microglia, astrocytes, OPCs, inhibitory neurons, and excitatory neurons. We used three different approaches to train the aging clock models for each cell type. In the first approach, we used the log-normalized gene expression values of individual cells, while in the second approach, we used the average of the log-normalized gene expression across all cells from a given donor and cell type, resulting in cell-type-specific simple pseudobulk samples (Fig. 3a and Supplementary Data 5a-l). Further, given that the averaging in the simple pseudobulk approach would result in only one data point per donor per cell type, thereby reducing variance, we used a third approach similar to what was described in Buckley et al 2023^4^. We randomly sampled (i.e. bootstrapped) a fixed number of cells and averaged the expressions into cell-type-specific bootstrapped pseudobulk samples (Fig. 3a), resulting in multiple data points per donor per cell type (Supplementary Data 5m-r).

**Figure 3.**
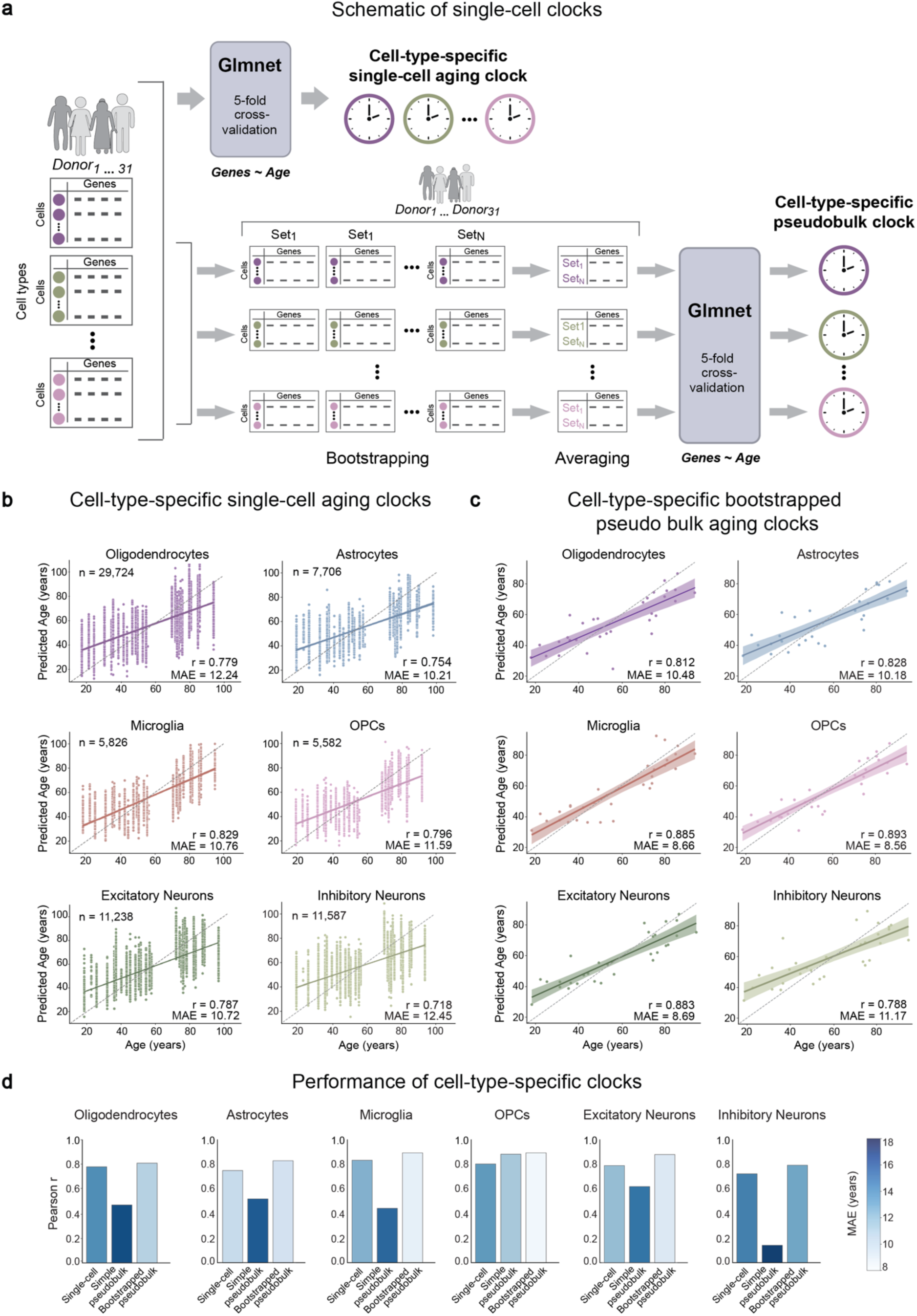
Development and evaluation of cell-type-specific transcriptomic aging clocks for the human brain. (a) Schematic representation of the different approaches in the development of the aging clocks using the ElasticNet regression model via the GlmNet algorithm. Cell-type-specific single-cell aging clocks were trained and tested directly on the (log-normalised) single-cell gene expression values, while the cell-type-specific (simple or bootstrapped) pseudobulk aging clocks were trained and tested on aggregated gene expression data of cells derived from a given cell type (b-c) Relationship between chronological age and the age predicted from the test rounds of (b) cell-type-specific single-cell aging clocks and (c) cell-type-specific bootstrapped pseudobulk clocks. (d) Bar plots showing Pearson’s correlation coefficients and mean absolute errors (MAE, represented by the intensity of blue colour in the bars) of each approach in each cell type. All correlation tests were performed using the stats.pearsonr function in SciPy, with significance based on a p-value < 0.05.

We found that the cell-type-specific single-cell aging clocks showed a statistically significant positive correlation between the predicted age and the chronological age for all cell types examined, with correlation coefficients 0.74 – 0.83 and mean absolute errors (MAE) of 10.2 – 12.2 years (Fig. 3b, Supplementary Data 6a-f and 7). The age prediction performance improved when we tested the cell-type-specific bootstrapped pseudobulk aging clocks (correlations 0.78 – 0.89, and MAEs 8.5 – 11.2 years) (Fig. 3c, Supplementary Data 6m-r and 7). However, the cell-type-specific simple pseudobulk aging clocks, with the exception of OPCs, performed relatively poorly with correlation coefficients as low as 0.4 and MAE as high as 18 years in oligodendrocytes (Fig. 3d, Supplementary Data 6g-l and 7), suggesting that the bootstrapped pseudobulk and the single-cell approaches captured more variation, allowing for greater accuracy in age predictions.

Next, we evaluated non-cell-type-specific pseudobulk aging clocks at the neuron level (based on aggregated gene expression data of inhibitory neurons and excitatory neurons), glia level (based on aggregated gene expression data of oligodendrocytes, astrocytes and OPCs) and across cells of all major cell types (based on aggregated gene expression) (Supplementary Fig. 3a and Supplementary Data 6s-aa). The bootstrapped-pseudobulk clocks in all three cases made predictions with high positive correlation, with a correlation coefficient of 0.9 and MAE of 8 years in the all-cells pseudobulk clock for example (Supplementary Fig. 3b, Supplementary Data 6s-aa and 7). Like the cell-type-specific case, we observed that the bootstrapped pseudobulk aging clocks showed better performance in age prediction compared to the simple pseudobulk clocks. For instance, the glia-level simple pseudobulk clock had a correlation coefficient of 0.2 while in the bootstrapped versions the correlation coefficient was around 0.8 (Supplementary Fig. 3b, Supplementary Data 6s-aa and 7). This further suggests that bootstrapping prior to training leads to more accurate predictions, possibly by sampling a greater range of variation in the data. Additionally, the similar performance in the bootstrapped neuron and glia level clocks suggest that the age-related transcriptomic changes can also be modelled across cells in the neuronal lineage and in the glial lineage. These results highlight the importance of capturing dynamic differences between cell types, which non-cell-type-specific clocks, despite their high performance, fail to address.

In summary, the overall performance of the cell-type-specific aging-clocks indicates that human brain aging can be accurately defined in each of the major cell types at the single-cell level, based on single-nuclei transcriptomic data from human post mortem prefrontal cortex tissue. In addition, these results further confirm the differences in age-related dynamics between cell types in the brain and the importance of studying aging at a cell-type-specific level.

### Validation of cell-type-specific aging clocks on independent single RNA sequencing datasets

To test the applicability of the aging clock models on independent datasets, we selected publicly available snRNA-seq datasets containing neurotypical adult human post mortem prefrontal cortex cohorts from two recent studies that have a broad and continuous age range and come from different subregions of the prefrontal cortex^18,32^ (Supplementary Table 1). In the Fröhlich et al dataset^18^, we used 33 control samples from the orbitofrontal cortex, which ranged in age from 26 to 84 years with a median PMI of 29.75 (6.5 to 50) hours (Supplementary Table 1). From the Velmeshev et al. dataset^32^, we used 12 adult frontal (10) or cerebral cortex (2) samples, aged 19 to 54 years, with a median PMI of 16.5 (6 to 27) hours (Supplementary Table 1). In all cases, we focused on the major cell types as used above. In addition, to verify whether the datasets were comparable, we performed label transfer and projected the UMAP structure using common anchors between the training and the independent datasets (Supplementary Fig. 4). We found that the clusters corresponding to various cell types in the UMAPs of both Fröhlich et al^18^ (Supplementary Fig. 4a) and Velmeshev et al^32^ (Supplementary Fig. 4b) aligned with that of the training dataset. Further, the predicted labels had high prediction scores for the matching original labels of all the cell types (Supplementary Fig. 4c,d, Supplementary Data 8a and c), with 90.5% matches in the Frohlich et al dataset^18^ (Supplementary Data 8b), and 98% matches in the Velmeshev et al dataset^32^ (Supplementary Data 8d), suggesting that majority of the cells were comparable.

We applied the trained clocks to the different datasets and checked the correlation of the predictions with the chronological age of the donors using Pearson’s correlation coefficient and the mean absolute error (MAE) of the predictions. In the Fröhlich et al datasets^18^, all clocks showed a statistically significant positive correlation between the predicted age and the chronological age. The level of correlation varied between the different cell types, with excitatory neurons showing the highest correlation and inhibitory neurons showing the lowest (Fig. 4a, b, c, Supplementary Data 9 and 11). The cell-type-specific single-cell aging clocks predicted with low correlation coefficients between 0.22 and 0.54 and MAE between 7.6 and 10 years (Fig. 4a, c, Supplementary Data 9a-f and 11), while the cell-type-specific bootstrapped pseudobulk aging clocks predicted with higher correlation coefficients between 0.56 and 0.78 and MAE between 7.1 and 15.6 years (Fig. 4b, c, Supplementary Data 9m-r and 11). In the cohort of Velmeshev et al^32^, the cell-type-specific single-cell aging clocks showed statistically significant, but generally low, correlations between 0.15 and 0.3 (Supplementary Figure 5a, c, Supplementary Data 10a-f and 11), whereas the predictions of the cell-type-specific bootstrapped pseudobulk aging clocks showed no statistically significant correlation in any of the cell types except inhibitory neurons and OPCs (Supplementary Fig. 5b, c, Supplementary Data 10m-r and 11).

**Figure 4.**
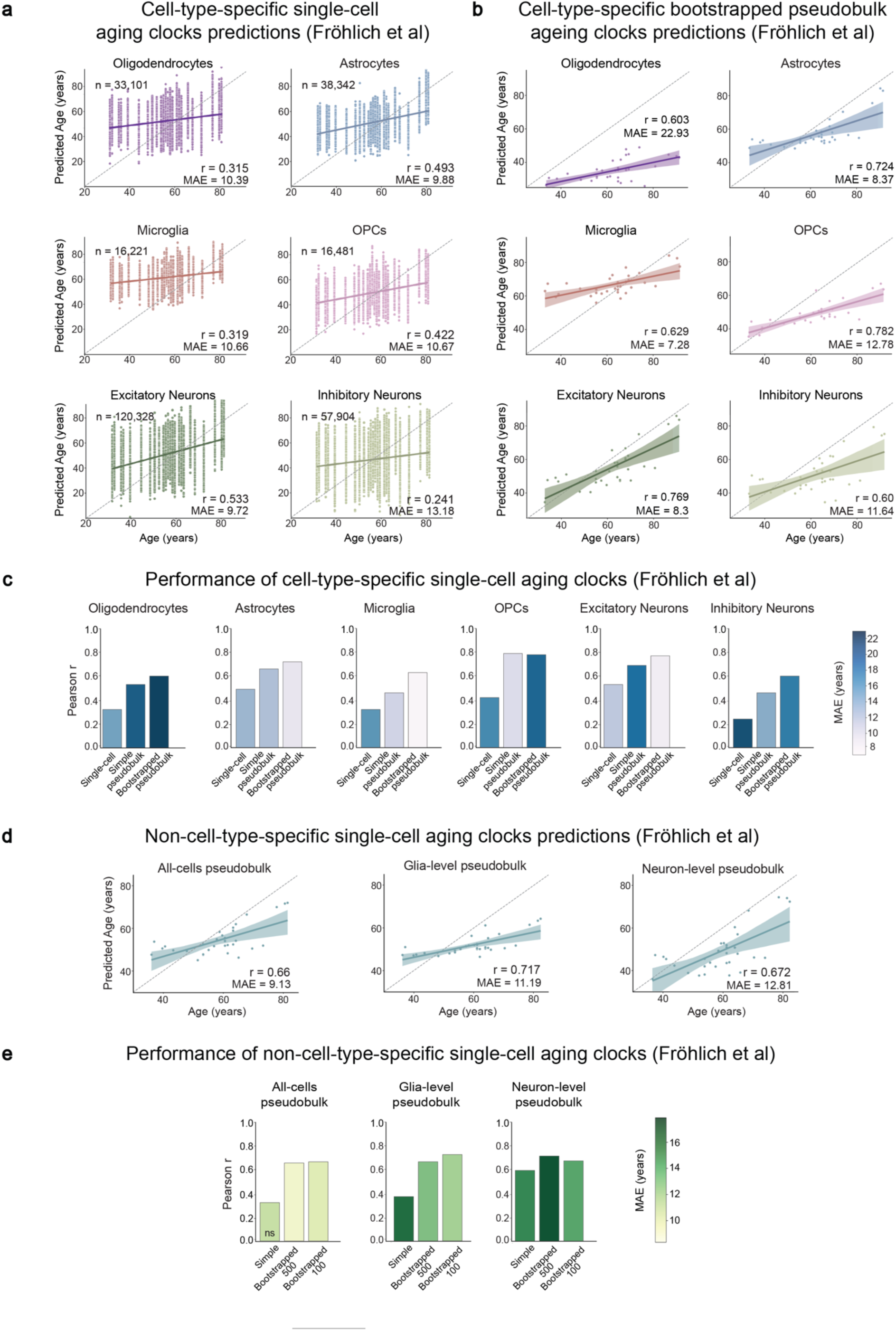
Validation of aging clocks on Fröhlich et al. dataset^18^. All plots show results from the validation of various aging clocks in Fröhlich et al dataset^18^. (a-b) Relationship between chronological age and predicted age in each cell, upon using (a) the cell-type-specific single-cell aging clock, and (b) the cell-type-specific bootstrapped-pseudobulk aging clock approaches. (c) Bar plots showing Pearson’s correlation coefficients and mean absolute errors (MAE, represented by the intensity of blue colour in the bars) of each of the cell-type-specific approaches in each cell type. (d) Relationship between the chronological age and the predicted age based upon using the different non-cell-type-specific pseudobulk clock approaches. (e) Bar plots showing Pearson’s correlation coefficients and mean absolute errors (MAE, represented by the intensity of green colour in the bars) of each of the non-cell-type-specific approaches. All correlation tests were performed using stats.pearsonr function of SciPy with significance based on a p-value < 0.05.

The non-cell-type-specific bootstrapped-pseudobulk aging clocks (for all cells, neuron and glia level), showed overall high positive correlations (r = 0.64 – 0.71) in the Fröhlich cohort (Fig. 4d, e, Supplementary Data 9s-y and 11). In the Velmeshev cohort on the other hand, interestingly, none except the neuron-level clock showed a statistically significant correlation (Supplementary Fig. 5d, e, Supplementary Data 10s-y and 11).

In summary, we found that the cell-type-specific bootstrapped-pseudobulk clocks showed a higher correlation and lower MAE in comparison to the single-cell clocks, possibly due to lower transcriptional noise. While the differences in the results between the datasets indicate dataset-specific effects on clock performance, we can conclude that the bootstrapped-pseudobulk clocks can predict biological age in the independent datasets, at a cell-type-specific level.

## DISCUSSION

In this study, we have generated a high-quality single-nucleus RNA sequencing dataset of post mortem human brains from a variety of prefrontal cortex subregions, with short PMIs and a broad age range spanning all three age groups of adulthood. The dataset has been used to develop human single cell-based transcriptomic aging clocks for each of the major cell types in the human prefrontal cortex, which can closely predict age at a cell-type specific level. The clocks predict age in all the major cell types, with the best performance in the bootstrapped pseudobulk aging clocks, where for example the OPCs, microglia and excitatory neuron clocks have correlation coefficients of around 0.9, and the single cell clocks of the same cell types with almost equal accuracy, with correlation coefficients around 0.8 in the test samples of the training dataset.

Epigenetic clocks have previously been applied to the brain^15,33–37^ and make predictions with high accuracy. However, these clocks do not capture the cell-type-specific information that is important for the brain, given its wide range of cell types^38^ and cell-type-specific vulnerabilities^39^. While cell type-specific epigenetic clocks were developed by using bulk brain tissue, sorted brain cells and deconvolution algorithms, this method has limitations (e.g. lower cell-type resolution) compared to the analysis of directly measured single-cells ^40^. Although single-cell DNA methylation approaches exist and single-cell DNA methylation clocks have been developed for other organ systems^10,11^, high cost, high sparsity, low-throughput, and low and random coverage in combination with the binary nature of the data pose major limitations for the development of aging clocks and their applicability^41–43^. In this context, single-nucleus RNA sequencing offers higher coverage, is scalable and high-throughput at a lower cost^44^, and its quantitative nature makes it more feasible for the development of cell-type-specific aging clocks. Overall, the use of cell-type-specific transcriptomic clocks are more feasible given the high prevalence of single-nucleus transcriptomics in the study of human brain aging and neurodegenerative diseases.

It is important to note that while both single-cell and bootstrapped-pseudobulk clocks showed high correlations in the training dataset, their performance in independent datasets was lower and varied. For example, the Velmeshev et al^32^ post mortem samples originated from across the frontal cortex and parts of cerebral cortex, which differ from the prefrontal cortex. This regional difference can introduce an increased variability in cell specific transcriptomic aging profiles. Additionally, the dataset included several middle-aged samples, where the direction of changes can be more complex when compared to the remaining age groups^45^. Overall, the differences in performance may be due to variations in sample preparation method, tissue quality, PMI, quality control thresholds, brain regional specificity and sequencing. Establishing thresholds for the minimum number of genes or cells required for optimal clock accuracy may help to better understand such differences. In addition, the donors in the training dataset were of limited geographical origin, so the variations were under-represented. Although increasing the sample size with equal representation of different ethnicities and sexes may help to mitigate the above issues, the availability of good quality post mortem tissue in adulthood in such large numbers is a major challenge. The single-cell clocks showed particularly poor performance, whereas the bootstrapped-pseudobulk clocks showed much better performance, for example, excitatory neuron and OPC clocks made predictions with correlation coefficients close to 0.8. This falls in line with previous observations in Buckley et al 2023^4^, and suggests that aggregating and bootstrapping the data may have overcome the transcriptomic noise associated with single-nucleus RNA sequencing. There is an urgent need to set standards and define the minimum criterion for further cell-type specific single cell transcriptomic clock development. In summary, we describe here a human brain aging snRNAseq dataset that has allowed us to construct transcriptomic clocks for all major cell types of the human brain. In age-related neurodegenerative diseases, such as Alzheimer’s disease, several age-related changes are often accelerated^46–50^. The aging clocks presented here can potentially help predict the acceleration of aging and reveal the selective vulnerability of key cell types. The next step is to further develop aging clocks so that they can be applied to samples obtained from late-onset neurodegenerative diseases such as Alzheimer’s and Parkinson’s disease. Moreover, it is important to develop additional region-specific clocks and subtype specific cell clocks to shed light on how aging affects cellular variability and vulnerability of the human brain.

## METHODS

### Human Post mortem Prefrontal Cortex Tissue

Human post mortem brain tissue samples (Table 1) were obtained from the Human Brain Tissue Bank (Semmelweis University, Budapest, Hungary). The tissue samples were collected from 31 individuals who were not diagnosed with any psychiatric disorder and neurodegenerative disorders. The brains were removed with a PMI of 2 – 12 hours, drop frozen using dry ice, and stored at −80 °C until further dissection. Samples were punched out from the lateral surface of the prefrontal cortices of the diseased subjects using specialized microdissection needles with 8- and 15-mm internal diameters. While most of the samples were dissected from the ventrolateral prefrontal cortex, some originated from the dorsolateral prefrontal cortex and from the middle frontal gyrus (the dorsal part of which belongs to the dorsolateral prefrontal cortex and its ventral part to the ventrolateral prefrontal cortex). The dissected cortical tissue pellets included both gray and white matter portions within the gyrus. Samples were collected in 1.5 mL Eppendorf tubes and stored at −80 °C until further use. Throughout the microdissection procedure, the tissue samples were kept frozen.

### Single-Nuclei Isolation

The nuclei isolation from the frozen post mortem brain tissue was performed as described previously^51,52^. Briefly, the tissue was thawed and dissociated in ice-cold lysis buffer [0.32 M sucrose, 5 mM CaCl_2_, 3 mM MgAc, 0.1 mM EDTA, 10 mM tris-HCl (pH 8.0), and 1 mM dithiothreitol] using a 1 ml tissue douncer (Wheaton). The homogenate was carefully layered on top of a sucrose cushion [1.8 M sucrose, 3 mM MgAc, 10 mM tris-HCl (pH 8.0), and 1 mM dithiothreitol] before centrifugation at 30,000 *g* for 2 hours and 15 minutes. After supernatant removal, the pelleted nuclei were softened for 10 minutes in 50 μl of nuclear storage buffer [15% sucrose, 10 mM tris-HCl (pH 7.2), 70 mM KCl, and 2 mM MgCl_2_] before being resuspended in 300 μl of dilution buffer [10 mM tris-HCl (pH 7.2), 70 mM KCl, and 2 mM MgCl_2_] and filtered through a cell strainer (70 μm). Nuclei were stained with Draq7 and run through FACS (FACS Aria, BD Biosciences) at 4°C at a low flow rate using a 100 μm nozzle (reanalysis showed >99% purity). 8,500 nuclei were sorted and used for downstream applications (snRNA Sequencing). Detailed protocol can be found at doi: dx.doi.org/10.17504/protocols.io.5jyl8j678g2w/v1.

### Single-nucleus RNA Sequencing

Nuclei intended for single-nucleus RNA-sequencing (8,500 nuclei per sample) were directly loaded onto the Chromium Next GEM Chip G or Chromium Next GEM Chip K Single Cell Kit along with the reverse transcription master mix following the manufacturer’s protocol for the Chromium Next GEM single cell 3′ kit (PN-1000268, 10x Genomics), to generate single-cell gel beads in emulsion. cDNA amplification was done as per the guidelines from 10x Genomics using 13 cycles of amplification. Sequencing libraries were generated with unique dual indices (TT set A) and pooled for sequencing on a Novaseq6000 using a 100-cycle kit and 28-10-10-90 reads.

### Single-nucleus RNA Sequencing Analysis

Raw base calls were demultiplexed to obtain sample-specific FastQ files and reads were aligned to the GRCh38 genome assembly using the *Cell Ranger* pipeline (10x Genomics Cell Ranger 7.0.0; RRID:SCR_017344)^44^ with default parameters and include-introns set to true. The resulting matrix files were used for further analysis.

All downstream analysis was done using *R* (v4.3.1; RRID:SCR_001905) and the standard workflow of *Seurat* (v4.3.0; RRID:SCR_016341)^53^. Nuclei with read counts between 1,200 and 100,000, gene counts between 800 and 12,000, and less than 5% mitochondrial transcripts were alone included for the analysis. The data was integrated using *Harmony* (v0.1.1; RRID:SCR_022206)^54^, where both sequencing batches and individual samples were regressed out, and the first 39 principal components were chosen. Clusters resolved at a resolution of 0.2 were annotated based on canonical cell type markers of the human prefrontal cortex. Further, any doublets or multiplets detected using *scDblFinder* (v1.13.10; RRID:SCR_022700)^55^ were removed. Differential gene expression analysis was done using the *FindMarkers* function and the Wilcoxon Rank Sum Test, between different age groups, namely, old vs young, old vs middle-age and middle-age vs young. Genes with adjusted p-value less than 0.05 and absolute log two-fold-change greater than 0.5 were considered significant. Mitochondrial genes and sex genes were excluded from further downstream analysis. Enrichment of GO terms corresponding to significant differentially expressed genes were tested with gene over-representation test using *ClusterProfiler* package (v4.8.2; RRID:SCR_016884), with Bonferroni-Hochberg correction for multiple testing.

### Aging clock models

Chronological age was predicted based on the log-normalized gene expression values using the Python implementation of the Glmnet algorithm for the ElasticNet regression model^30,31^. Mitochondrial and sex-related genes were removed from the data resulting in 35,578 genes in total to use as features. Separate models were trained directly on the cell data of the six major cell types yielding cell-type-specific single-cell age prediction models (i.e., aging clock) for oligodendrocytes, astrocytes, microglia, OPCs, excitatory neurons and inhibitory neurons. To prevent information leakage between training and test data, we trained and tested the age prediction models with donor-level 5-fold cross-validation (i.e. in one iteration, we trained a model using cells of the 80% of the donors and predicted the age of cells of the remaining 20% of the donors). We experimented with different alpha parameter values (alpha = 0, 0.5, and 1) and selected the models trained with alpha = 0.5 due to their best overall performance (also considering external performance) of predicting the age of single-cells. Performance was measured by Pearson’s correlation coefficient (r) between the chronological and predicted age, as well as by mean absolute error (MAE) of the predictions (Supplementary Data 3.).

Beside the cell-type-specific single-cell aging clock approach, we also developed cell-type-specific pseudobulk aging clocks using generated pseudobulk samples in two ways: (i) simple pseudobulk where gene expressions were averaged over all cells of a given cell type of a given donor, resulting in one pseudobulk sample for each donor per and cell-type; and (ii) bootstrapped pseudobulk where 100 samples were generated for each donor and cell-type by randomly sampling and averaging a given number of cells (for oligodendrocytes: 200, astrocytes: 50, microglia: 50, OPCs: 50, excitatory neurons: 100, and inhibitory neurons: 100). Then, similarly to the single-cell approach, separate models were trained for each cell-type with donor-level 5-fold cross-validation, and with different values for parameter alpha (alpha = 0, 0.5, and 1). The predicted age of a donor given by the cell-type-specific bootstrapped pseudobulk aging clocks was calculated by the average prediction of the donor’s 100 pseudobulk samples for the given cell-type.

We also developed non-cell-type-specific pseudobulk aging clocks at the glia-level, neuron-level, and for all cells (simple and bootstrapped). The glia-level clocks were based on oligodendrocytes, astrocytes and OPCs; the neuron-level clocks were based on excitatory and inhibitory neurons; and the all-cells clocks used all cells from the six major cell types. We tested both simple and bootstrapped pseudobulk approaches for the three levels. For the simple pseudobulk approach, we generated samples by averaging the gene expressions over all cells of the given cell types (neuron, glia, and all-cells) of a donor. For the bootstrapped approach, 100 samples per donor were generated by taking the average over *k* randomly sampled cells of the given level. We tested the approach with k = 500 and 100 cells. In total, we tested nine types of non-cell-type-specific aging clocks (glia-level/neuron-level/all-cells by simple, bootstrapped with 100 cells, and bootstrapped with 500 cells approach). The training and validation of the clocks were performed the same way as described previously^12^.

### Label transfer and UMAP projection

Using *Seurat* (v4.4.0; RRID:SCR_016341)^53^ and *R* (v4.3.1; RRID:SCR_001905), Seurat’s recommended workflow for label transfer and UMAP projection was used. The processed single-nuclei RNA sequencing data of the training dataset and the external datasets were subset to include only the major cell types. Additionally, in the Velmeshev et. al. dataset, samples corresponding to adulthood, over 18 years of age, and originating from the frontal or the cerebral cortex were alone included. Given that the Fröhlich et. al. dataset was originally normalized by Pearson’s residuals, the raw counts were log-normalized to keep them comparable to the training dataset. Further, top 2,000 highly variable genes in the Fröhlich et. al. dataset were identified using the *FindVariableFeatures* function, followed by re-computing PCs and UMAP embeddings, using the *RunPCA* and *RunUMAP* functions, respectively.

With default parameters, *FindTransferAnchors* function with PCA as reference reduction was used to find common anchors, where the training dataset with first 18 PCs was used as reference while the Fröhlich et. al. or Velmeshev et. al. datasets were used as query. Using the derived anchors, *TransferData* function was used to predict cell type annotations and the *MapQuery* function was used to project UMAP embeddings in the query datasets.

### Aging clock validation on independent external datasets

To examine the generalizability of the proposed aging clocks, we applied them to two independent external datasets. A subset of the snRNA-seq data of Velmeshev et al.^32^ namely, the frontal cortex samples of adult donors (age > 18 years) were used for external validation of the clocks, as well as orbitofrontal cortex snRNA-seq data of the control samples of Fröhlich et al.^18^. In both cases, the processed gene expression data was utilised, and further processing of the samples was performed as described above in the section ‘Aging clock models’. Missing values were imputed by the average expressions of the missing genes of the training dataset, where the average was calculated on the samples the clocks were trained on (e.g. on the bootstrapped samples for the bootstrapped clocks). Due to the 5-fold cross-validation described above, five regression models were generated for each clock. All five models were applied to each external sample and their average prediction was used for evaluation. The evaluation of the clocks on the independent datasets was done as described in ‘Aging clock models’.

## DATA AND CODE AVAILABILITY

All data needed to evaluate the conclusions in the paper are present in the paper and/or the Supplementary Materials section. The training dataset will be deposited in the Gene Expression Omnibus (GEO). In case of validation datasets, the processed snRNA-seq data of Velmeshev et al.^32^ can be accessed at https://cellxgene.cziscience.com/collections/bacccb91-066d-4453-b70e-59de0b4598cd, while that of Fröhlich et al.^18^ is available under GEO superseries GSE254569. Clock models have been added to Supplementary Data 5. The code for the single-nuclei RNA sequencing analysis is available at https://github.com/chanmur/ppfctx_aging_snRNAseq_analysis_2025, and that for the aging clock part is at https://github.com/SZTAKI-SU-Rejuvenation-Group/ppfctx_aging_snRNAseq_clock_2025.

## ACKNOWLEDGEMENTS

We are thankful to all members of the HCEMM-SU Neurobiology and neurodegenerative diseases research group, the Laboratory of Molecular Neurogenetics, HUN-REN-SZTAKI-SU Rejuvenation research group and the Human Brain Bank. We thank the Strategic Research Area’s MultiPark (Multidisciplinary Research on Parkinson’s Disease) and the Lund Stem Cell Center for infrastructure support.

## AUTHOR CONTRIBUTIONS

M.P. and É.R. provided the microdissected human brain samples used in the study. A.A., J.G.J. and D.A.M.A. carried out the single nuclei sequencing experiments under the supervision of J.J. A.A.A. designed and visualized the figures. C.M. designed the single-cell computational framework and analysed the data under the supervision of K.P., Y.S. and J.J. Y.S. and R.G. verified the analytical methods. E.Z.-P. developed and applied the single-cell transcriptomic clocks under the supervision of C.K. C.M. wrote the manuscript with support from K.P., C.K. and J.J. and with input from all authors. K.P. and C.K. conceived the study and were in charge of overall direction and planning. K.P., J.J. and C.K. supervised the project. All authors provided critical feedback and helped shape the research, analysis and manuscript.

## ETHICS STATEMENT

This study adhered to the principles outlined in the Declaration of Helsinki and the Ethical Rules for Using Human Tissues for Medical Research in Budapest, Hungary (HM 34/1999). Activity of Human Brain Tissue Bank, Semmelweis University has been authorised by the Committee of Science and Research of Ethic of Ministry of Health, Hungary (189/KO/02.6008/2002/ETT) and the Regional Committee of Science and Research Ethics (No.32/1992/TUKEB). All experiments involving human post mortem samples described in this study were conducted under the ethical approval number (IV/2627-1 /2021/EKU).

## SUPPLEMENTARY INFORMATION

### Supplementary Table

**Supplementary Table 1.**
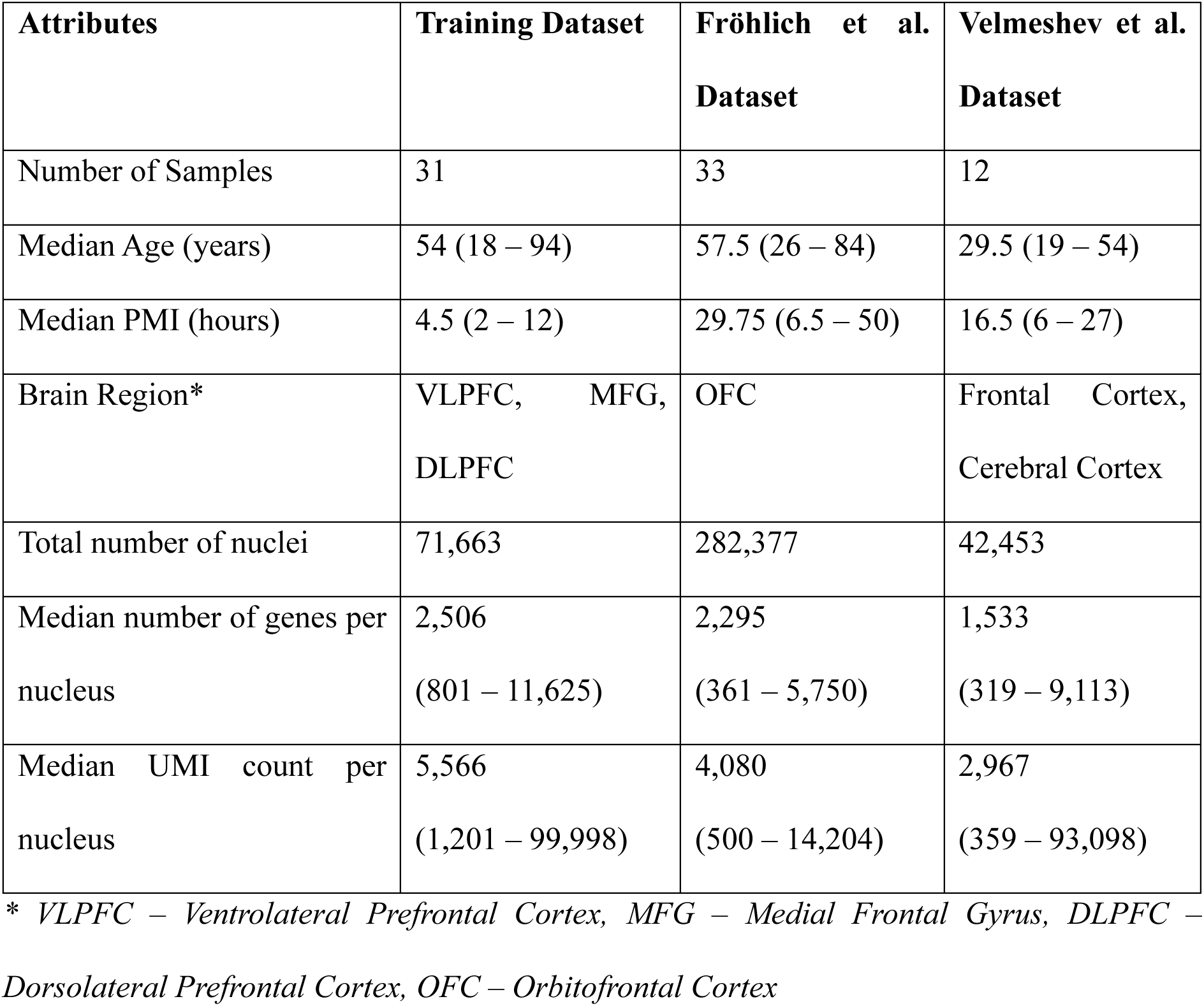
Summary of sample details in the training and external datasets for major cell types after QC.

| Attributes | Training Dataset | Fröhlich et al.<br>Dataset | Velmeshev et al.<br>Dataset |
| --- | --- | --- | --- |
| Number of Samples | 31 | 33 | 12 |
| Median Age (years) | 54 (18 – 94) | 57.5 (26 – 84) | 29.5 (19 – 54) |
| Median PMI (hours) | 4.5 (2 – 12) | 29.75 (6.5 – 50) | 16.5 (6 – 27) |
| Brain Region* | VLPFC, MFG,<br>DLPFC | OFC | Frontal Cortex,<br>Cerebral Cortex |
| Total number of nuclei | 71,663 | 282,377 | 42,453 |
| Median number of genes per nucleus | 2,506<br>(801 – 11,625) | 2,295<br>(361 – 5,750) | 1,533<br>(319 – 9,113) |
| Median UMI count per nucleus | 5,566<br>(1,201 – 99,998) | 4,080<br>(500 – 14,204) | 2,967<br>(359 – 93,098) |

\* *VLPFC – Ventrolateral Prefrontal Cortex, MFG – Medial Frontal Gyrus, DLPFC – Dorsolateral Prefrontal Cortex, OFC – Orbitofrontal Cortex*

### Supplementary Figures

**Supplementary Figure 1.**
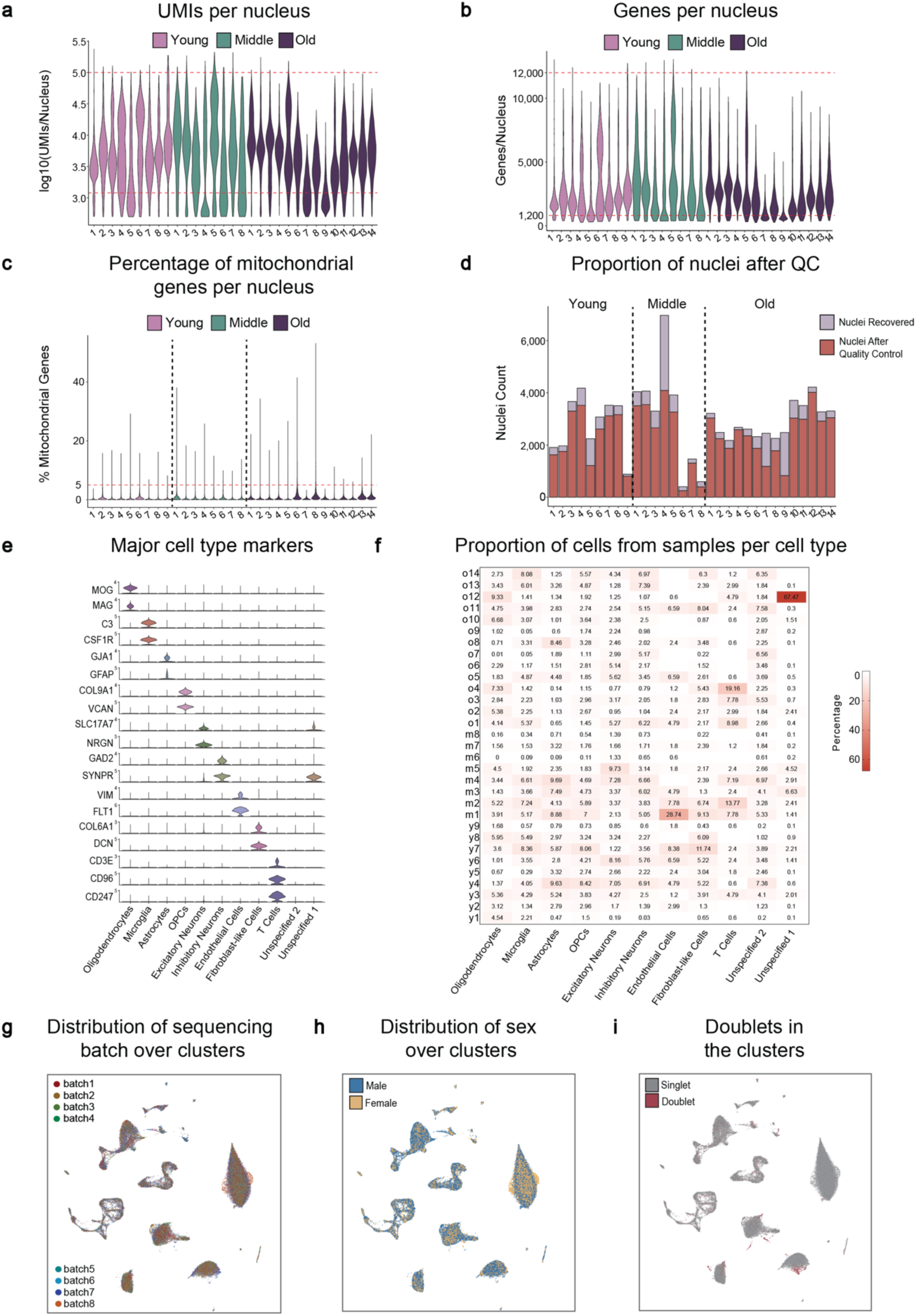
Quality and cell composition of the dataset. Related to Figure 1. (a) Violin plot showing the distribution of number of UMIs per nucleus per sample in log10 scale. (b) Violin plot showing the distribution of number of genes per nucleus per sample. (c) Violin plot showing the distribution of percentage of mitochondrial genes per nucleus. (d) Bar plot showing the proportion of nuclei per sample after quality control. In a-d, the upper and lower limit of the range included are indicated by the red dashed lines. (e) Violin plots showing the expression of additional canonical markers corresponding to cell types in the prefrontal cortex. (f) Table showing the proportion of cells per sample in each cell type. (g) UMAP plot with annotation of sequencing batch in every cluster. (h) UMAP plot with annotation of sex in every cluster. (i) UMAP plot showing annotation of predicted singlets and doublets.

**Supplementary Figure 2.**
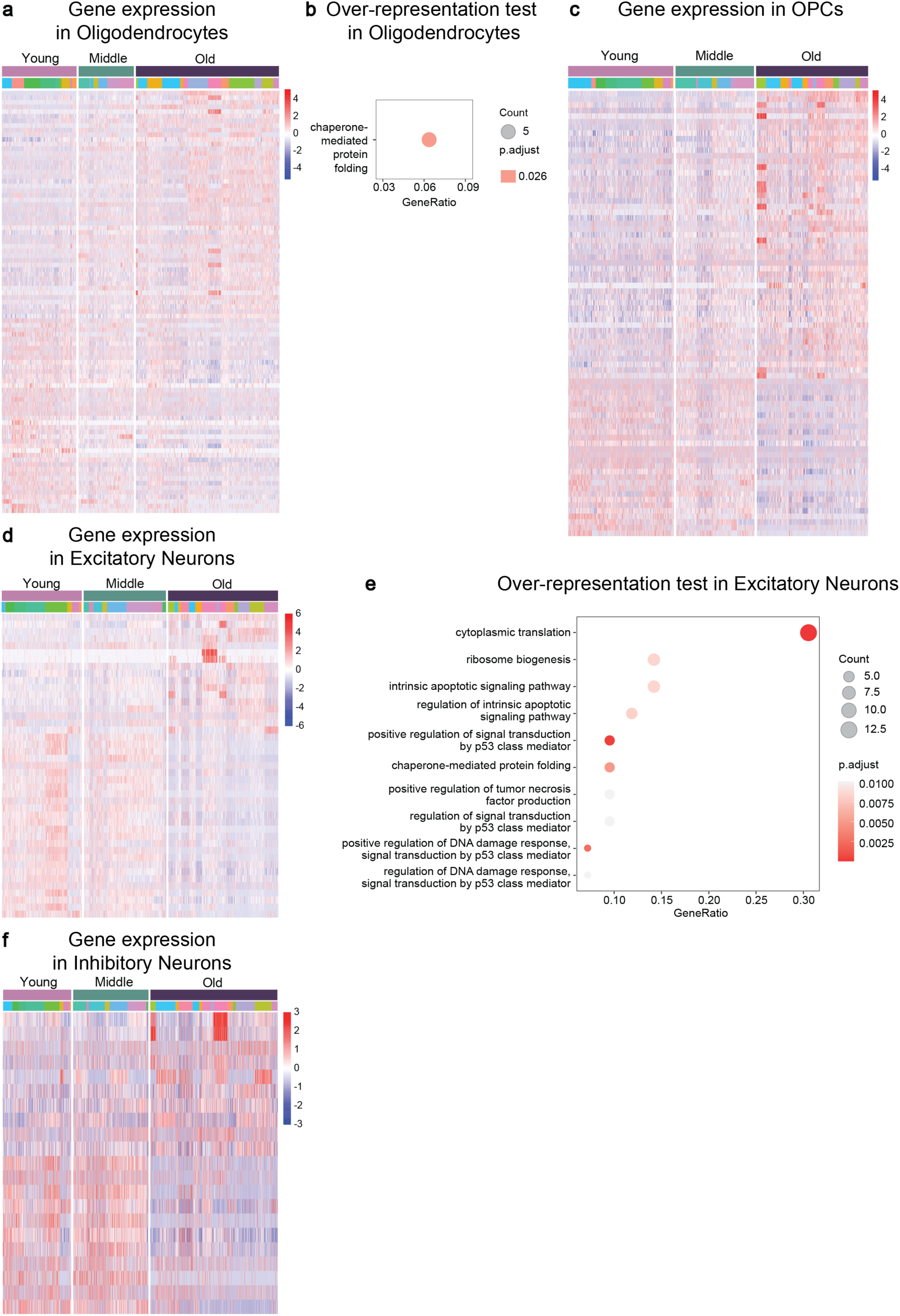
Age-related changes in various major cell types. Related to Figure 2. (a) Heatmap showing the expression of significantly differentially expressed genes in oligodendrocytes when comparing old vs young age groups. (b) Plot showing gene ontology terms enriched from genes differentially expressed in old oligodendrocytes. (c) Heatmap showing the expression of significantly differentially expressed genes in OPCs when comparing old vs young age groups. (d) Heatmap showing the expression of significantly differentially expressed genes in excitatory neurons when comparing old vs young age groups. (e) Plot showing gene ontology terms enriched from genes differentially expressed in old excitatory neurons. (f) Heatmap showing the expression of significantly differentially expressed genes in inhibitory neurons when comparing old vs young age groups. *p < 0.05; ClusterProfiler’s inbuilt test for gene-overrepresentation analysis along with Bonferroni-Hochberg correction for multiplicity was used in b and e. Different colours below the young, middle and old ages bar represents cells originating from different individuals in a, c, d and f.

**Supplementary Figure 3.**
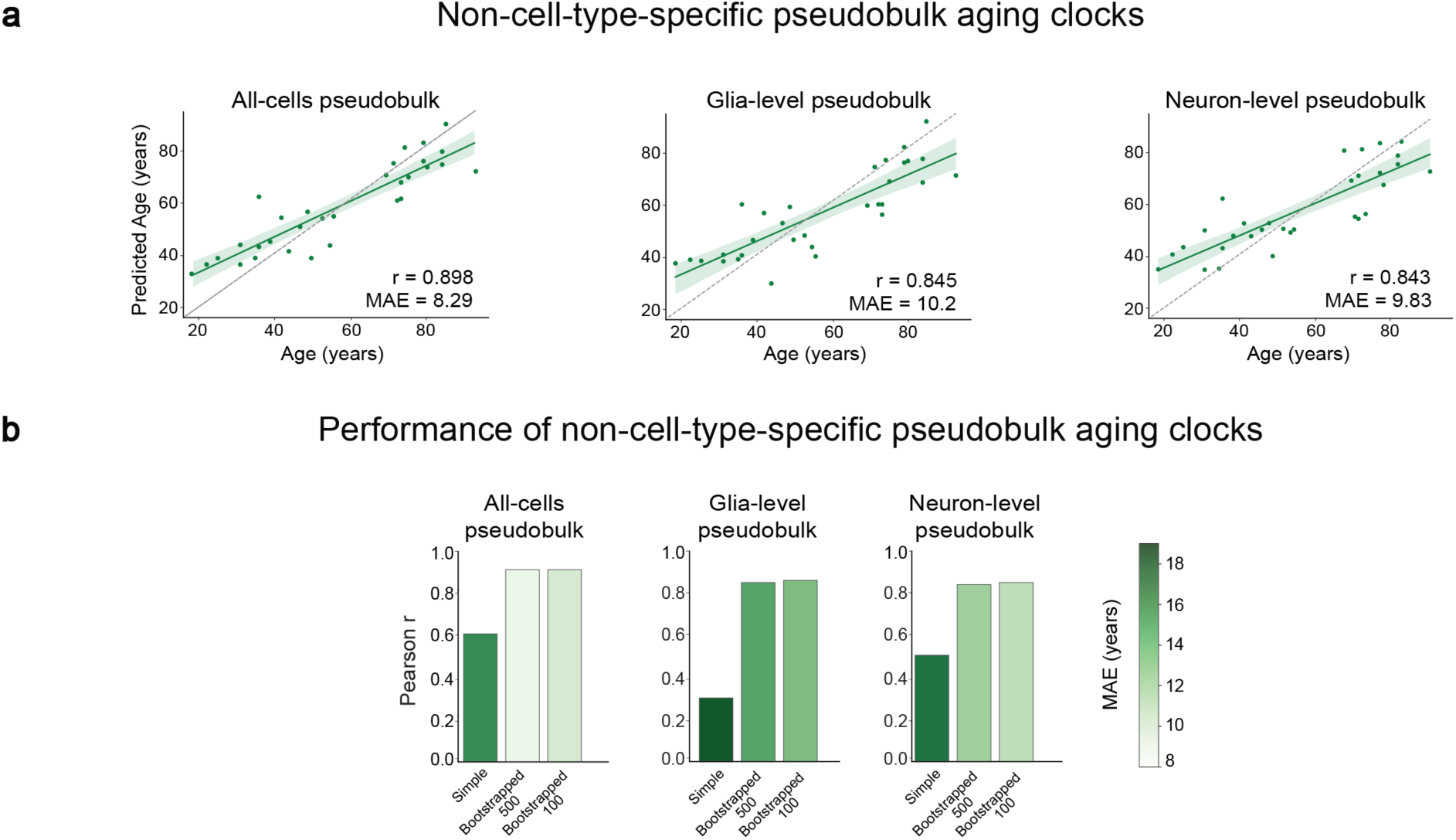
Development and evaluation of non-cell-type-specific pseudobulk aging clocks. (a) Relationship between the chronological age and the predicted age by using bootstrapped pseudobulk aging clock approaches trained on all-cells, at the glia-level, and neuron-level, respectively. (b) Pearson r and mean absolute error (MAE) from each pseudobulk clock approach. For each level (all-cells/glia-level/neuron-level), three versions are displayed: simple, bootstrapped with 500 cells, and bootstrapped with 100 cells, respectively. All correlation tests were performed using the stats.pearsonr function of SciPy with significance based on a False Discovery Rate (FDR) < 0.05.

**Supplementary Figure 4.**
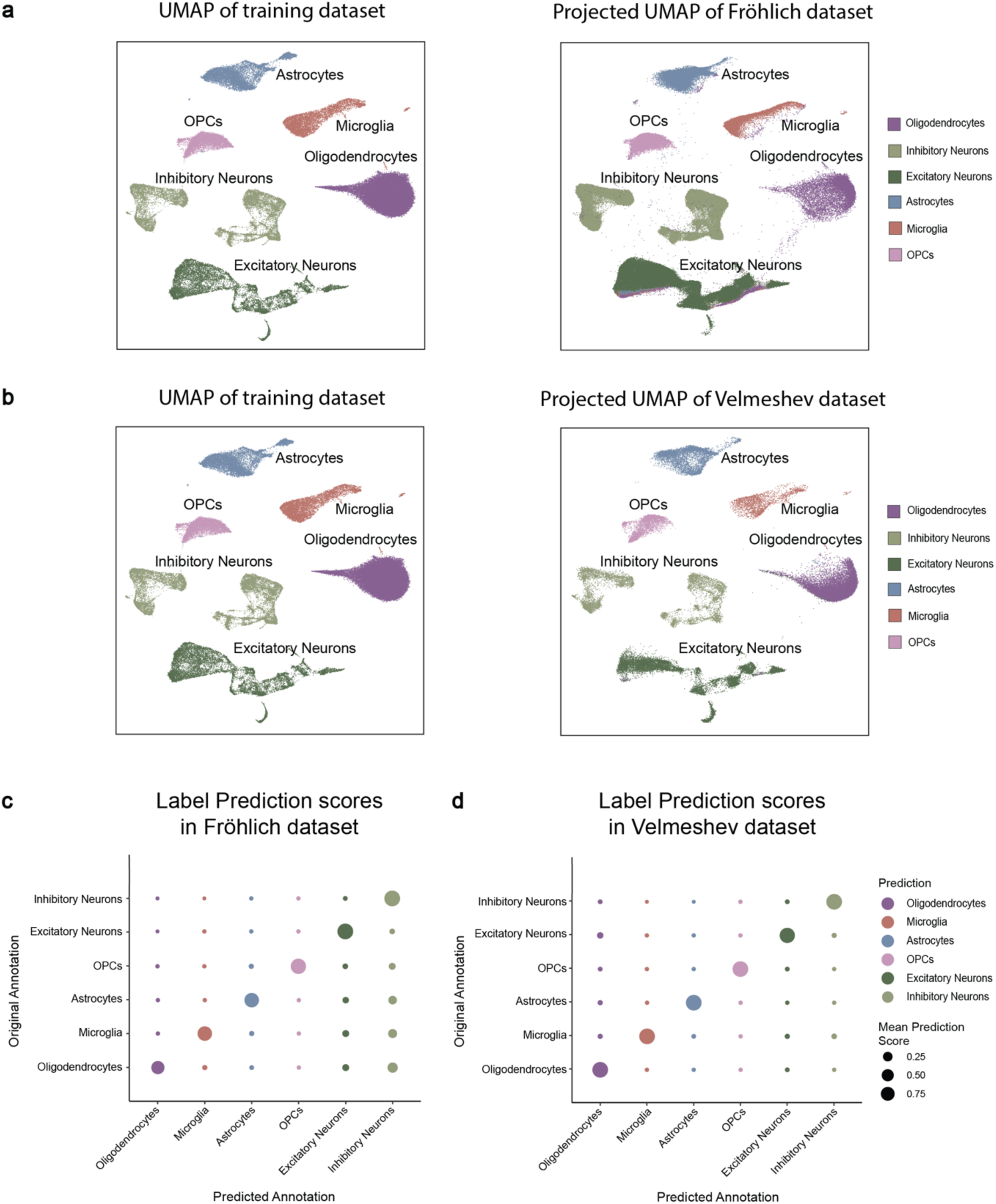
Comparison of cell type annotation and clustering between the training dataset and the validation datasets. Related to Figure 4. (a) UMAPs of training dataset as reference and its unimodal projection on Fröhlich et al^18^ dataset. (b) UMAPs of training dataset as reference and its unimodal projection on the Velmeshev et al^32^ dataset. (c) Bubble plot showing the mean prediction score for different combinations of original annotations and predictions in the Fröhlich et al^18^ dataset with the training dataset as reference. (d) Bubble plot showing the mean prediction score for different combinations of original annotations and predictions in Velmeshev et al^32^ dataset with training dataset as reference.

**Supplementary Figure 5.**
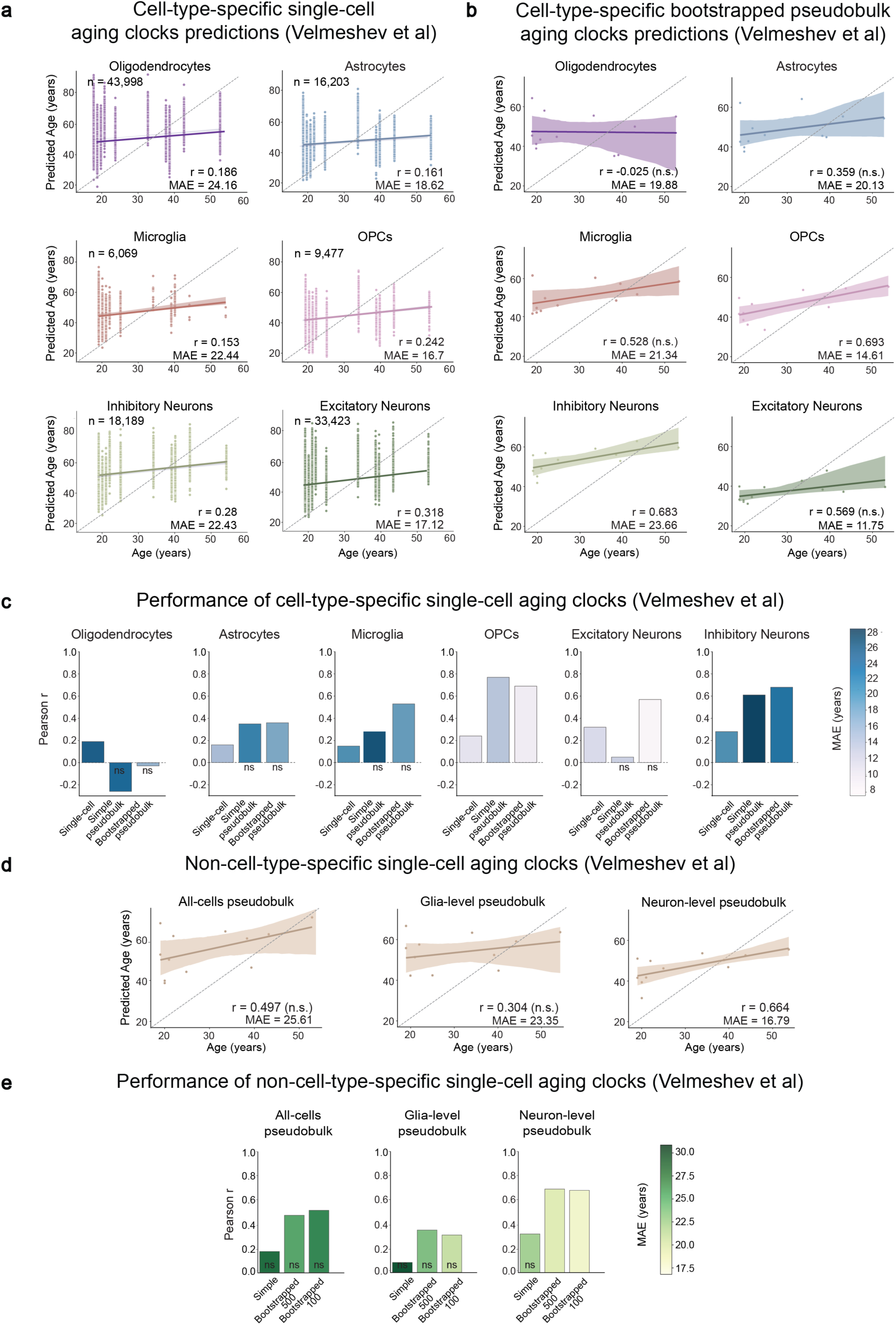
Validation of aging clocks on Velmeshev et al. dataset^32^. All plots show results from the validation of various aging clocks in Velmeshev et al dataset^32^. (a-b) Relationship between chronological age and predicted age in each cell, upon using (a) the cell-type-specific single-cell aging clock, and (b) the cell-type-specific bootstrapped-pseudobulk aging clock approaches. (c) Bar plots showing Pearson’s correlation coefficients and mean absolute errors (MAE, represented by the intensity of blue colour in the bars) of each of the cell-type-specific approaches in each cell type. (d) Relationship between the chronological age and the predicted age based upon using the different non-cell-type-specific pseudobulk clock approaches. (e) Bar plots showing Pearson’s correlation coefficients and mean absolute errors (MAE, represented by the intensity of green colour in the bars) of each of the non-cell-type-specific approaches. All correlation tests were performed using stats.pearsonr function of SciPy with significance based on a p-value < 0.05.

### List of Supplementary Data

**Supplementary Data 1:** Sample information and distribution of cells from different cell types.

**Supplementary Data 2:** Gene expression in each cell type in the training dataset.

**Supplementary Data 3:** Differential gene expression analysis results from all major cell Types across all age-group comparisons.

**Supplementary Data 4:** Gene over-representation test results from all comparisons.

**Supplementary Data 5:** Aging clocks: regression models of each of the 27 aging clocks.

**Supplementary Data 6:** Predicted age from the testing rounds of each of the 27 aging clocks.

**Supplementary Data 7:** Summary of aging clock performances in the training dataset.

**Supplementary Data 8:** Label transfer prediction results in the external datasets.

**Supplementary Data 9:** Predicted age given by the 27 aging clocks in Fröhlich et. al. dataset.

**Supplementary Data 10:** Predicted age given by the 27 aging clocks in Velmeshev et. al. dataset.

**Supplementary Data 11:** Summary of aging clock performances in the external datasets.

